# Megakaryocyte activation and mobilization from the bone marrow in response to trauma and hemorrhagic shock in mice

**DOI:** 10.64898/2026.02.10.704825

**Authors:** Isabelle C Becker, Eleanor J Smith, Nicola Dark, Jordi L Tremoleda, Harriet E Allan, Paul Vulliamy

## Abstract

Severe injuries result in acute changes in platelet number and function, but the impact of trauma and hemorrhagic shock on megakaryocytes (MKs) in the initial hours after injury have not been studied in detail. Using a murine model of trauma-hemorrhage, we identified rapid changes in MK morphology and mobilization into bone marrow sinusoids, changes that were detectable within one hour. Levels of several alpha-granule derived proteins were elevated in the bone marrow, and co-culture of naive MKs with bone marrow supernatant from injured mice resulted in similar changes to those observed in the model. These results illustrate that trauma-hemorrhage results in a hyperacute alteration in the bone marrow micro-environment that alters MK activity within an hour of injury.

## MAIN TEXT

Tissue injury and hemorrhage represent an immediate challenge to the survival of living organisms. A fundamental aspect of any successful response to injury and blood loss initially involves achieving hemostasis at sites of bleeding, followed by a process of wound healing and tissue repair.^1^ These responses require maintenance of a circulating pool of blood cells, which in turn entails an adaptable hematopoietic system that can modulate its output during times of acute physiological stress. However, a detailed understanding of how hematopoiesis is affected during the critical early window after injury is currently lacking. This is an important knowledge gap, because maladaptive or dysregulated responses to severe trauma occur frequently and are associated with adverse clinical outcomes.^2^

Platelets are pivotal both to an effective hemostatic response and in orchestrating subsequent inflammation and tissue repair.^3^ We and others have previously described a range of alterations in platelet number and function that occur within the first hours of severe injury in humans, but the acute impact of trauma and hemorrhage on megakaryocytes (MKs) has not previously been studied.^4,5^ Much of our understanding of how thrombopoiesis is regulated involves pathways that take several days (for example, thrombopoietin-induced megakaryopoiesis and subsequent platelet production), which are not relevant to the immediate post-injury response.^6^ Increasing evidence supports the concept that MKs can respond to stress, i.e. during infection, an emergency hematopoiesis from hematopoietic stem cells (HSCs) can occur in order to bypass slower pathways of platelet production that predominate in homeostasis.^7,8^ However, it is not known how and whether MKs can respond within minutes (as opposed to hours-days) of a physiological stressor, as is needed in the setting of severe trauma and blood loss.

To investigate how traumatic injury affects the MK lineage, we used an established mouse model of trauma and hemorrhagic shock (THS) which closely recapitulates the acute coagulopathy and organ dysfunction observed in injured humans.^9,10^ As previously described,^9,10^ THS mice underwent tissue injury followed by 60 minutes of hemorrhagic shock (target mean arterial pressure 25-35 mmHg), after which we harvested the bone marrow for further processing. Sham mice underwent anesthesia and arterial cannulation only (**supplemental figure 1A**). Compared to sham mice, THS mice had significantly lower blood pressure during the shock phase, and higher terminal serum lactate (**supplemental figure 1B-C**). To investigate MK characteristics and behavior in situ within the bone marrow, we performed confocal microscopy of whole femur cryosections (supplemental methods). Compared to sham mice, we found a reduction in the number of MKs present within the bone marrow in THS mice (**figure 1A**) and a reduction in cell size (**figure 1B**), a feature of less mature MKs.^11^ In addition, MKs in THS mice had reduced circularity and solidity (**figure 1C-D**), morphological features that are characteristic of migratory cells.^11,12^ Overall cellularity in the bone marrow was reduced in THS compared to sham mice (**figure 1E**), consistent with previous reports in injured humans.^13^ These findings prompted us to assess the localization of MKs within the bone marrow of THS mice in more detail. We indeed observed a significantly greater proportion of MKs in close contact with and within sinusoids compared to sham mice (**figure 1F-G)**, suggesting an egress of mature MKs into the bloodstream upon traumatic injury. Supporting the indications of altered MK maturity in the bone marrow, examination of ploidy distribution revealed that in THS mice there was a shift towards less mature MKs shown by an increase in 8N cells but a reduction in 16N cells (**figure 1H**). Collectively this suggests active MK mobilization towards and into the vasculature in the hyperacute post-injury period.

**Figure 1:**
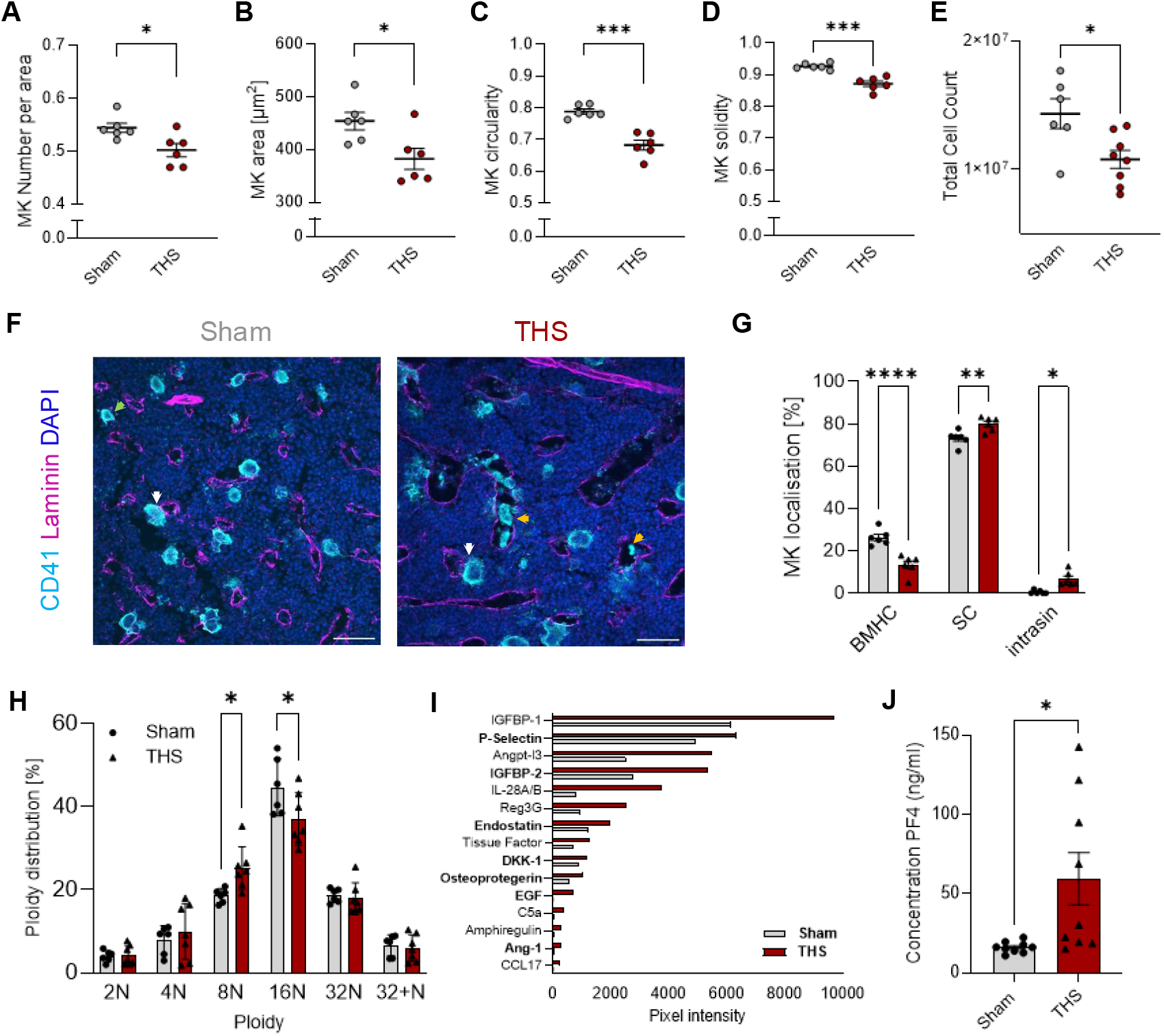
Characteristics of megakaryocytes in the bone marrow after trauma hemorrhage. Confocal microscopy was performed to determine **(A)** megakaryocyte (MK) numbers, **(B)** MK area; **(C)** MK circularity and **(D)** MK solidity in sham (grey) and THS mice (red), n=6 mice. **(E)** Quantification of total bone marrow cellularity in sham (grey) and THS (red) mice, n=6-8. **(F)** Representative confocal images of bone marrow cryosections from sham and THS mice stained for CD41 (cyan), laminin (magenta) and DAPI (blue). Scale bars: 50 µm. MKs within the bone marrow haematopoietic (BMHC) are indicated by green arrows, MKs with sinusoidal contact (SC) are indicated by the white arrows, and intra-sinusoidal (intrasin) MKs are indicated by orange arrows (Zeiss LSM880, 20x objective). **(G)** Quantification of MK localisation within the BMHC, with SC or intrasin in sham (grey circles) and THS (red, triangles) mice, n=6 mice. **(H)** Flow cytometric analysis of ploidy distribution (propidium iodide) of MKs isolated from the bone marrow of sham (grey circles) and THS (red triangles) mice. Analysis of cytokine profiles within the bone marrow supernatant of sham and THS mice showing **(I)** significantly upregulated cytokines in the bone marrow supernatant (BMS) of THS mice compared to sham mice (BMS pooled from 8 mice per condition) ranked by highest abundancy, with MK-derived cytokines highlighted in bold. **(J)** Concentration of Platelet Factor 4 (PF4) in BMS from sham and THS mice measured using a PF4 ELISA, n=8 mice. Analysis was performed using an unpaired t-test; * p<0.05. Analysis was performed using an unpaired t-test or Two-Way ANOVA with Šídák’s multiple comparisons test; * p<0.05, **p<0.01, *** p<0.0005, **** p<0.0001.

To determine whether injury-induced changes within the bone marrow microenvironment might contribute towards these changes, we measured the levels of a broad range of cytokines in the bone marrow supernatant (BMS). We identified an upregulation of 15 cytokines in the BMS of THS mice, however, none of the upregulated cytokines have previously been implicated in cell migration. In contrast, the majority of upregulated cytokines are of megakaryocytic origin and stored within their α-granules, including P-selectin, EGF, IGFBP-2, Ang-1 and DKK-7 (highlighted in bold; **figure 1I and supplemental figure 2**) suggesting an increase in granule secretion from MKs upon injury. To confirm this, we measured levels of the canonical α-granule protein platelet factor 4 (PF4) in BMS and found a significant increase in concentration in THS mice compared to sham mice (**figure 1J**). Interestingly, only 4 out of the 15 proteins upregulated in the bone marrow were also elevated in blood plasma of THS mice (**supplemental figure 2**), indicating a distinct post-injury microenvironment within the bone marrow compared to circulation. Of note, the majority of upregulated plasma cytokines were either involved in mediating platelet activation (including CCL21, CX3CL1, Resistin), or directly derived from platelet α-granules (including PF4, Ang-1, GDF-15, CXCL5, MMP3 and MMP9; **supplemental figure 2**), in line with previous studies identifying increased platelet activation in trauma patients.^4^

To investigate how upregulated bone marrow cytokines may affect subsequent megakaryo- and thrombopoiesis, we next performed a series of co-culture experiments using HSCs and MKs derived from wild-type mice incubated with the BMS from sham and THS mice (supplemental methods).^13^ HSCs treated with BMS from THS mice generated smaller MKs compared to HSCs treated with sham BMS (**figure 2A**), while there was no difference in the number of MKs (**figure 2B**). In line with the smaller size, we also observed a shift in ploidy distribution towards lower ploidy following exposure to THS BMS (**figure 2C**). Finally, mature MKs treated with BMS derived from sham or THS mice produced greater numbers of proplatelets (**figure 2D)**, which were larger in area compared to MKs treated with BMS from sham mice (**figure 2E-F**).

**Figure 2:**
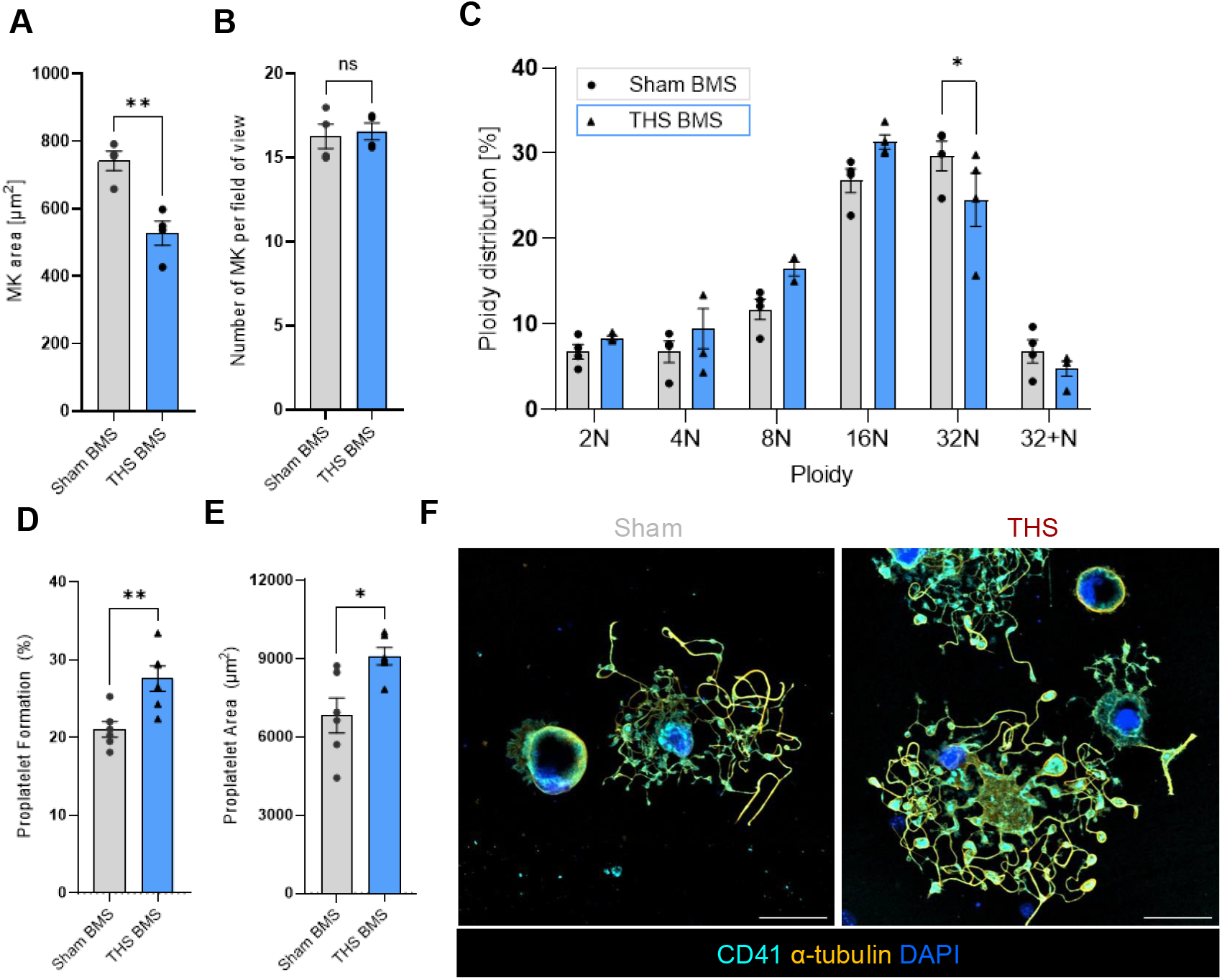
Characterisation of the effects of sham and THS bone marrow supernatant on megakaryocyte maturation and proplatelet formation. Quantification of **(A)** megakaryocyte (MK) area and **(B)** MK number following incubation of haematopoietic stem and progenitor cells (HSPCs) with thrombopoietin (50 ng/ml) and recombinant hirudin (100 U/ml) and bone marrow supernatant (BMS) from sham or THS mice for 3 days, n=4. **(C)** Flow cytometric analysis of MK ploidy distribution (propidium iodide) following incubation with sham (grey) and THS (blue) BMS, n=4. Proplatelet formation was assessed by confocal microscopy following incubation of mature MKs with sham or THS BMS for 24 hours. Quantification of **(D)** the percentage of proplatelet-forming MKs and **(E)** proplatelet area of sham and THS BMS treated MKs, n=6. **(F)** Representative confocal microscopy (Nikon AXR; 40x Objective, 1.6xZoom) images showing proplatelet formation of sham and THS BMS-treated MKs stained with CD41 (cyan), α-tubulin (yellow), and DAPI (blue). Scale bars: 50 µm. Analysis was preformed using ImageJ. Analysis was performed using a paired t-test or a Two-Way ANOVA with Šídák’s multiple comparisons test; * p<0.05, **p<0.01.

We and others have previously identified early changes in platelet number and function during THS that correlate with worse outcomes in patients.^4^ Whether altered MK differentiation and platelet production may counteract increased platelet consumption during THS has not been investigated to date. The present findings represent the first description of how trauma-hemorrhage affects the form and functional properties of bone marrow-resident MKs within the ‘golden hour’ after injury. We found evidence of immediate mobilization of mature MKs towards and into the vasculature, resulting in reduced MK counts and overall ploidy. Moreover, the significant increase in MK-derived growth factors and cytokines in the BMS strongly implies rapid activation of and enhanced granule secretion from bone marrow-resident MKs. We were further able to identify that an upregulation of these cytokines affects subsequent megakaryo- and thrombopoiesis in vitro, resulting in the generation of lower ploidy MKs but enhanced proplatelet formation. Our findings in this murine model are consistent with historical reports of reduced bone marrow cellularity in injured humans, while mobilization of MKs, despite being a feature of other pathologies such as cancer, has not been previously observed in THS.^14,15^ Although our findings emphasize an important, immediate response of MKs towards traumatic injury, the downstream effects on platelet production require further study, as do the mechanisms underpinning the changes we describe. Overall, our observations shed new light on the early post-injury bone marrow responses and demonstrate that thrombopoietic machinery can adapt within minutes of a physiological stressor.

