## Supplemental Methods & Figures for "Megakaryocyte activation and mobilization from the bone marrow in response to trauma and hemorrhagic shock in mice"

#### Mouse model of trauma and hemorrhagic shock

All studies were carried out under Project License PP6747232, approved by the university Animal Welfare and Ethical Review Body and UK Home Office, in compliance with the EU Directive 2010/63/EU. C57BL/6 mice (Charles River laboratories, Margate UK) aged 8-12 weeks, both male and female were anaesthetised with isofluorane. Cannulation of the right carotid artery was performed to allow for constant blood pressure monitoring and controlled haemorrhage. A severe blunt injury pattern was mimicked via closed bilateral tibial fractures using a haemostat followed immediately by midline laparotomy, with bilateral rectus muscle crushing using a haemostat. Blood was withdrawn via the carotid cannula to a target mean arterial pressure of 25-35 mmHg (Catpo SP 844, AD Instruments, UK) for 60 minutes. A terminal blood sample was collected via the carotid cannula, and the femora were extracted and placed in phosphate-buffered saline (PBS, D8537, Sigma Aldrich) or 4% paraformaldehyde (PFA, 043368-9M, Thermo Fisher Scientific) prior for further processing. Sham mice underwent anaesthesia and cannulation only, followed by terminal blood sample collection after 1 hour of anaesthesia and femora extraction. Serum Lactate was measured at the time of cannulation (baseline) and following terminal blood collection using an Accutrend Plus Meter (Roche).

#### Cryosectioning and immunofluorescence staining

Femora were isolated as described above and fixed using 4% PFA overnight. Afterwards, femora were moved into 10% sucrose/PBS (24h at 4°C), followed by 20% and 30% sucrose. Dehydrated femurs were embedded in a water-soluble embedding medium and frozen down at -20°C. Using a tape transfer system (Super Cryofilm Tape, C-FUF303, Section Lab), 10 µm sections were retrieved at a cryostat (Leica Biosystems). Sections were rehydrated in PBS for 20 min, non-specific binding was blocked using 10% donkey serum (D9663, Sigma Aldrich) and sections were stained with rat anti-mouse CD41 (MWReg30, MA5-16875, Invitrogen), rabbit anti-mouse laminin (L9393, Sigma-Aldrich) and 4′,6-diamidino-2-phenylindole (DAPI, Merck), followed by anti-rat Alexa Fluor 488 (A21208, Thermo Fisher Scientific) and anti-rabbit Alexa Fluor 647 (A31573, Invitrogen). Slides were washed in PBS containing 0.1% Triton X-100 (T8787, Sigma Aldrich) and mounted using Fluoroshield mounting media (F6182, Sigma Aldrich). Sections were imaged at a Zeiss LSM880 confocal microscope (20x objective). Whole femora were imaged and MKs were quantified using a TissueFax Plus imaging platform (10x objective). Analysis was performed using ImageJ (NIH).

**Isolation of bone marrow cells**

Following THS or sham procedures and a terminal blood collection, femora were collected and placed in PBS. Cells were isolated by centrifugation for 40 seconds at 2500 xg as previously described.[16] Total cell count was determined using a Countess II (Invitrogen).

**Analysis of ploidy distribution in total bone marrow cells**

Bone marrow cells were isolated as described above; red blood cells were lysed in ACK buffer (A1049201, Gibco) and cells were passed through a 70 µm cell strainer. Cells were centrifuged at 300 xg, then fixed and permeabilized in 70% ethanol/PBS for 30 min on ice. Cells were treated with RNAse A (EN0531, Thermo Fisher Scientific), stained with anti-CD41-FITC antibodies (133904, BioLegend) and propidium iodide (P4864 Sigma-Aldrich) for 30 min on ice, and ploidy distribution was assessed using flow cytometry (Cytek Aurora). Analysis was performed using FlowJo v.10.

**Bone marrow supernatant collection**

For the collection of the bone marrow supernatant (BMS), femora were isolated as described above. Bones were spun out into 100µL PBS, BMS was collected and frozen at -20°C.

**Blood plasma collection**

Following the terminal blood draw, peripheral blood plasma was mixed with 3.2% sodium citrate (BD, New Jersey USA) to a final concentration of 1:9. These samples were then centrifuged at 10,000 rpm for 10 minutes at room temperature. The supernatant plasma was then separated from the rest of the blood product, initially flash frozen in liquid nitrogen and then stored at -80°C.

**Cytokine array and enzyme-linked immunosorbent assay**

Cytokine levels were determined in BMS and blood plasma using a Proteome Profiler Mouse XL Cytokine Array (ARY028, R&D Systems), pooling samples from 8 mice per condition. Levels of platelet factor 4 within the BMS and blood plasma were determined using a mouse a mouse CXCL4/PF4 DuoSet ELISA Kit (DY595, BioTechne) according to the manufacturers' guidelines.

**Co-culture of bone marrow-derived haematopoietic stem cells with BMS**

Bone marrow cells were isolated from healthy C57BL/6 mice by centrifugation as described above. Cells were passed through a 70 μm cell strainer and incubated with a rat anti-mouse lineage panel (133307, BioLegend), followed by magnetic bead isolation using Dynabeads™ (11415D, Thermo Fisher Scientific). Lineage-depleted haematopoietic stem and progenitor cells (HSPCs) were cultured in DMEM containing 50 ng/ml thrombopoietin (TPO, 488-TO-005/CF, BioTechne), 100 anti-thrombin units (ATU) of recombinant hirudin (rHirudin; RE120A, Quadratech Diagnostics Ltd) and BMS from THS or sham mice (1:10 dilution) for days. Number and size of MKs were assessed by light microscopy (INCA6000), and ploidy distribution was assessed as described above. Analysis was performed using ImageJ (NIH) and FLowJo v.10.

**Co-culture of mature megakaryocytes with BMS**

Proplatelet formation in the presence of sham or THS BMS was assessed using mature MKs derived from in vitro differentiation of HSPCs. Mature MKs were enriched using size exclusion filtration[17], and proplatelet formation was determined in the presence of 100 ATU of rHirudin and BMS (1:10) from sham and THS for 24 hours on anti-mouse CD31-coated (102502, Biolegend) ibidi chambers. Proplatelets were stained with rat anti-mouse CD41 (MWReg30, MA5-16875, Invitrogen), mouse anti-α-tubulin (clone: B-5-1-2, T6074, Sigma Aldrich) and DAPI (Merck), followed by anti-rat Alexa Fluor 488 (A21208, Thermo Fisher Scientific) and anti-mouse Alexa

Fluor 555 (A-31570, Thermo Fisher Scientific). Proplatelets formation assessed using confocal microscopy (Nikon AXR; 40x objective). Analysis was performed using Image J (NIH).

**Supplemental Figure 1**

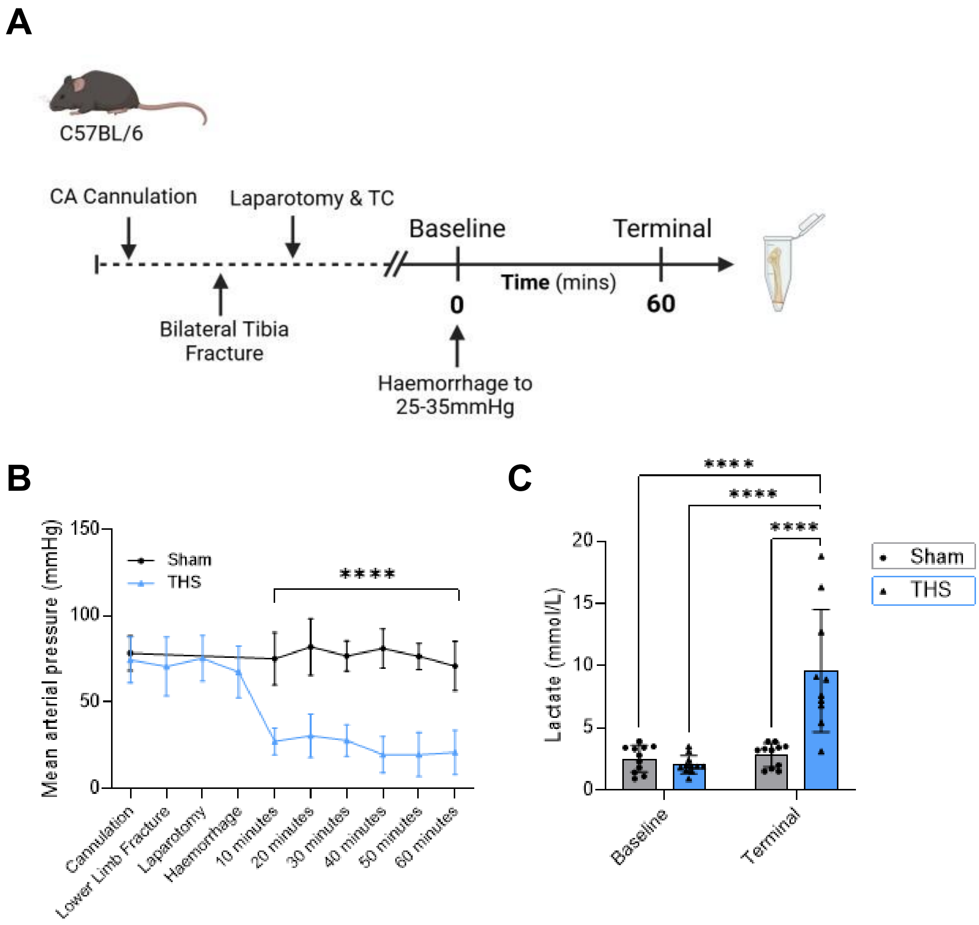


**Supplemental Figure 1: Experimental model of trauma and haemorrhagic shock. (A)** Procedures and timeline of interventions of trauma and haemorrhagic shock (THS) model. C57BL/6 mice aged 8-12 weeks, both male and female, were anaesthetised with isofluorane. Carotid (CA) cannulation was performed, followed by closed bilateral tibial fractures using a haemostat. Laparotomy and tissue crush (TC) was then performed using a haemostat to crush the rectus muscle bilaterally. Subsequently blood was withdrawn via the carotid cannula to a target mean arterial pressure of 25-25 mmHg for 60 minutes. A terminal blood sample was collected via the carotid cannula, and the femora were extracted and placed in phosphate-buffered saline or 4% paraformaldehyde prior to further processing. Sham mice underwent anaesthesia and cannulation only. **(B)** Mean arterial blood pressure measurements prior to and during haemorrhagic shock in sham and THS mice. **(C)** Serum lactate measurements at baseline (cannulation) and terminal timepoints in sham and THS mice. Analysis was performed using a Two-Way ANOVA with Tukey's multiple comparisons; ****p<0.0001.

### Supplemental Figure 2

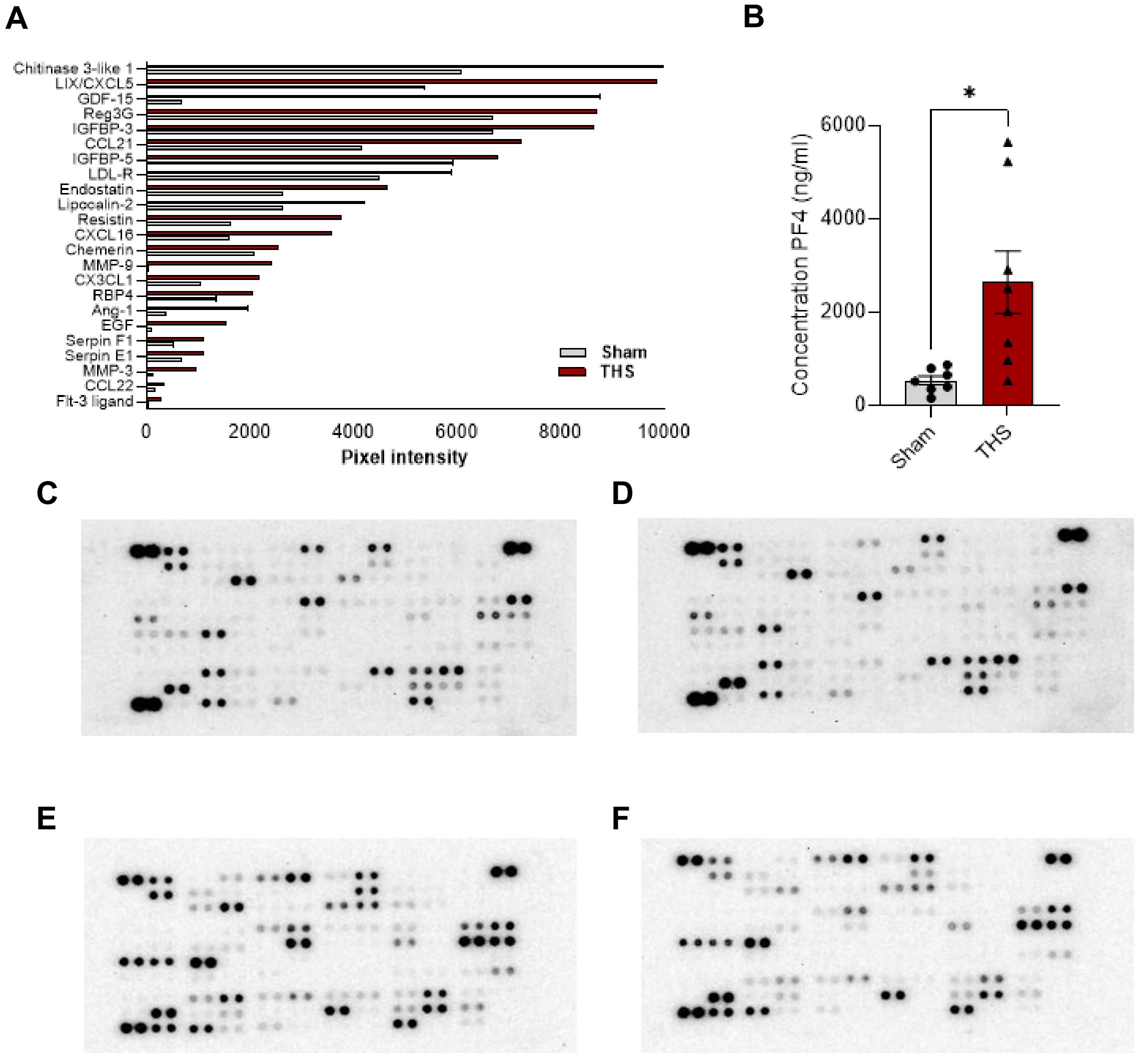


**Supplemental Figure 2**: **Cytokine array analysis and blots.** Analysis of cytokine profiles within the blood plasma of sham and THS mice showing **(A)** Significantly upregulated cytokines in blood plasma of THS mice compared to sham mice (blood plasma was pooled from 8 mice per condition). **(B)** Concentration of Platelet Factor 4 in the peripheral plasma in sham and THS mice measured using a PF4 ELISA. Analysis was performed using an unpaired t-test; * $p<0.05$. Proteome Profiler Mouse XL Cytokine Arrays were used to characterize bone marrow supernatant (BMS) and blood plasma composition from sham and THS mice. The blots for each condition are shown as follows **(C)** BMS, THS mice **(D)** BMS, sham mice **(E)** Plasma, THS mice **(F)** Plasma, sham mice.
